# Microbial ecology of acidic, biogenic gypsum: Community structure and distribution of extremophiles on freshly formed and relict sulfate deposits in a hydrogen sulfide-rich cave

**DOI:** 10.1101/2025.02.14.637391

**Authors:** Zoë E. Havlena, Katherine Lucero, Heather V. Graham, Jennifer C. Stern, Scott D. Wankel, Maurizio Mainiero, Daniel S. Jones

## Abstract

Sulfate minerals are abundant on the Martian surface, and many of these evaporite deposits are thought to have precipitated from acidic fluids. On Earth, gypsum (CaSO_4_•2H_2_O) and other sulfates sometimes form under acidic conditions, so exploring the extremophilic life that occurs in these mineral environments can help us evaluate the astrobiological potential of acid sulfate depositional settings. Here, we characterized the microbial communities associated with acidic gypsum deposits in a sulfuric acid cave, where sulfate precipitation is driven by sulfide-oxidizing bacteria and archaea. We used 16S rRNA gene sequencing and cell counts to characterize gypsum-associated microorganisms in freshly formed and relict deposits throughout the cave, in order to test hypotheses about how microbial community composition and abundance would vary with distance from the sulfidic water table and with the concentration of H_2_S(*g*) and other gases in the cave atmosphere. We found that actively-forming gypsum in the lower cave levels was colonized by low diversity communities of sulfide-oxidizing chemolithotrophs and other acidophiles that have few cells compared to other environments in the cave. The most abundant taxa were *Acidithiobacillus, Metallibacterium, Mycobacteria*, and three different *Thermoplasmatales*-group archaea, which occupied distinct niches based on proximity to sulfidic streams and the concentration of gases in the cave air. In contrast, deposits in older cave levels had more diverse communities that are dominated by chemoorganotrophic and methanotrophic taxa. These findings show that acidic sulfate deposits serve as habitats for extremophilic microorganisms, and broaden our knowledge of the life associated with terrestrial sulfates.

**Importance:** Gypsum and other sulfate salts are common on Mars, and many of these deposits are thought to have formed from acidic fluids early in the planet’s history. Understanding the life that survives and thrives in similar environments on Earth is therefore crucial for evaluating whether these Martian sulfates are or ever were habitable. One such environment where acidic gypsum occurs is in sulfuric acid caves, where extremophilic microorganisms drive the precipitation of sulfate minerals by oxidizing hydrogen sulfide gas from the cave atmosphere. Here, we characterized the communities of microorganisms on freshly formed and ancient gypsum in the Frasassi Caves, and found that the gypsum deposits hosted microbial communities that changed based on chemical energy availability and the age of the gypsum. Our findings underscore the importance of chemical and microbiological interactions in shaping habitable niches, and provide context for searching for past or present life in acidic Martian sulfates.

## Introduction

Gypsum (CaSO_4_•2H_2_O) and other sulfate minerals are prevalent on Mars, including strata at Gale crater that are being visited by the Curiosity Rover (1). Due to their association with aqueous conditions and extremophilic life on Earth, sulfate minerals represent potentially habitable environments in the past (2, 3), and these evaporite deposits are considered targets for life detection missions (4, 5). Many of the Martian hydrated sulfates are thought to have formed under acidic conditions (6, 7), and widespread evidence for acidic fluid alteration on Mars includes strata in Terra Sirenum’s Cross crater (8), the Meridiana Planum (9), and the Gale crater (10). On Earth, microbial life is abundant in extreme, gypsum-precipitating environments such as low pH playas and salars (11, 12), acid hot springs and volcanically-influenced sediments (13, 14), and acid mine drainage (15–17). In extreme environments, gypsum crystals can provide protection for endolithic microorganisms (18–21), and may possibly even be a source of bioavailable water (5). Acidic gypsum deposits can preserve dormant cells and fossil evidence for life (11, 22), perhaps even over long timescales (4). Earth’s acidic gypsum deposits are therefore valuable chemical analogues for Martian sulfates that can help understand how microorganisms use these mineral environments, and reveal the fate of biosignatures left behind in sulfates (4, 11, 23).

Acidic gypsum also forms in the terrestrial subsurface in sulfidic caves. Sulfidic caves develop through a process known as sulfuric acid speleogenesis (SAS), in which anoxic groundwaters with dissolved hydrogen sulfide (H_2_S) are exposed to oxygen at or near the cave water table. Sulfide-oxidizing microorganisms thrive at this mixing zone and catalyze the oxidation of H_2_S to elemental sulfur and sulfuric acid. This occurs both below the water table in cave streams and lakes, and above the water table on cave walls and ceilings where H_2_S degasses into the cave atmosphere (24, 25). Above the water table, microcrystalline gypsum forms as wall crusts where sulfuric acid corrodes subaerial limestone surfaces, in what is sometimes referred to as a “replacement solution” or “gypsum-replacement” process (SO_4_^2-^ + 2H^+^ + CaCO_3_ + H_2_O → CaSO_4_•2H_2_O + CO_2_). As SAS progresses, this secondary gypsum accumulates on cave wall and ceiling surfaces, and eventually sloughs off and accumulates as breakdown deposits on cave floors. Barring dissolution from contact with fresh surface drips or other cave waters, these “relict” gypsum deposits may persist for extended timescales, even up to 12 Ma in now-inactive SAS caves in New Mexico’s Guadalupe Mountains (26).

Italy’s Grotte di Frasassi (Frasassi Caves) are an active sulfidic cave system where these processes can be observed. SAS development occurs where H_2_S-rich groundwaters emerge at the modern water table, and new gypsum crusts precipitate near turbulent, degassing streams. Freshly-formed gypsum in the lower levels is extremely acidic (pH <2), and has a toothpaste-like microcrystalline texture, sometimes with larger selenite crystals (27–29). Immediately above degassing streams, it is often associated with extremely acidic microbial biofilms known as “snottites” (29–31), although gypsum crusts develop further from the zone where snottites are found (32, 33).

Relict gypsum deposits up to several meters thick can be found in older passages that are 10s to 100s of meters above the current water table (27). These areas have been progressively removed from the influence of the sulfidic aquifer by tectonic uplift and incision by the Sentino River, and form a series of sub-horizontal multi-story levels that record historic SAS processes dating back to the early and mid-Pleistocene (32, 34–36). Because these relict gypsum deposits are no longer exposed to degassing H_2_S, any microorganisms must rely on other and presumably more limited energy resources. Ancient SAS caves are generally oligotrophic (37), and energy sources for extant microbes colonizing speleothem and other mineral surfaces may include trace gases (38, 39), trace minerals in bedrock (40), and organic inputs (41).

Gypsum in active sulfuric acid caves like Frasassi may therefore offer unique insights into sulfate mineral-associated microbial communities in a subterranean environment where primary production is fueled by microbial chemolithoautotrophy. However, while the microbial communities that form acidic “snottite” biofilms have been well studied (e.g., (30)), little is known about the microbial community associated with the secondary gypsum itself.

Furthermore, we don’t know how long these deposits continue to serve as microbial habitats in relict sulfidic caves. We therefore sampled microbial communities associated with gypsum deposits throughout the Frasassi Caves, in order to (i) characterize microbial communities and processes in actively forming gypsum, and (2) evaluate if and how microbes inhabit older deposits. We hypothesized that communities on gypsum near the sulfidic water table would be dominated by taxa related to known sulfide-oxidizing chemolithotrophs and other acidophiles, and that these groups would gradually disappear further from the water table and be absent in relict gypsum from the older cave levels. We tested these hypotheses by using high-throughput 16S rRNA gene sequencing and cell counts to evaluate how microbial community composition and abundance varied with distance from the sulfidic water table and with the concentration of H_2_S(*g*) and other gases in the cave atmosphere.

## Results

### Sample collection and field observations

We sampled gypsum surfaces from multiple sites in the lower, actively forming levels, as well as from relict deposits in the older, upper levels (Table 1, Figs. 1 and 2). In the lower levels, we collected gypsum from three different passages with sulfidic streams: Ramo Sulfureo (RS), Pozzo dei Cristalli (PC), and Grotta Bella (GB). At these locations, we collected samples from areas immediately above streams where H_2_S(*g*) concentrations were as high as 33 parts per million by volume (ppmv), to areas more than 30 meters from the stream where H_2_S(*g*) was undetectable (Table 1). Gypsum crusts near sulfidic streams were consistently acidic (pH ≤2), and were usually damp toothpaste-like coatings on subaerial surfaces (Fig. 1a-c). At site RS, small crystals of elemental sulfur (S°) occurred with gypsum (Suppl. Fig. S1) We also collected some samples from the floor and wall along the entrance route in Grotta del Fiume (GdF) that leads to PC but is more than 25 meters from the nearest degassing stream, where H_2_S was not detectable by smell or portable gas sampling equipment (Table 1). Gypsum deposits in GdF and further from the stream at sites PC and GB were often dry and powdery, and pH could often not be measured in the field with pH strips. CO_2_ concentrations ranged from 800-5200 ppmv near streams to as low as 700 ppmv in the GdF passage, and other gases (NH_3_, SO_2_ and CO) were below detection. Complete sample information and associated geochemical data are provided in Supplementary Table S1.

**Figure 1.**
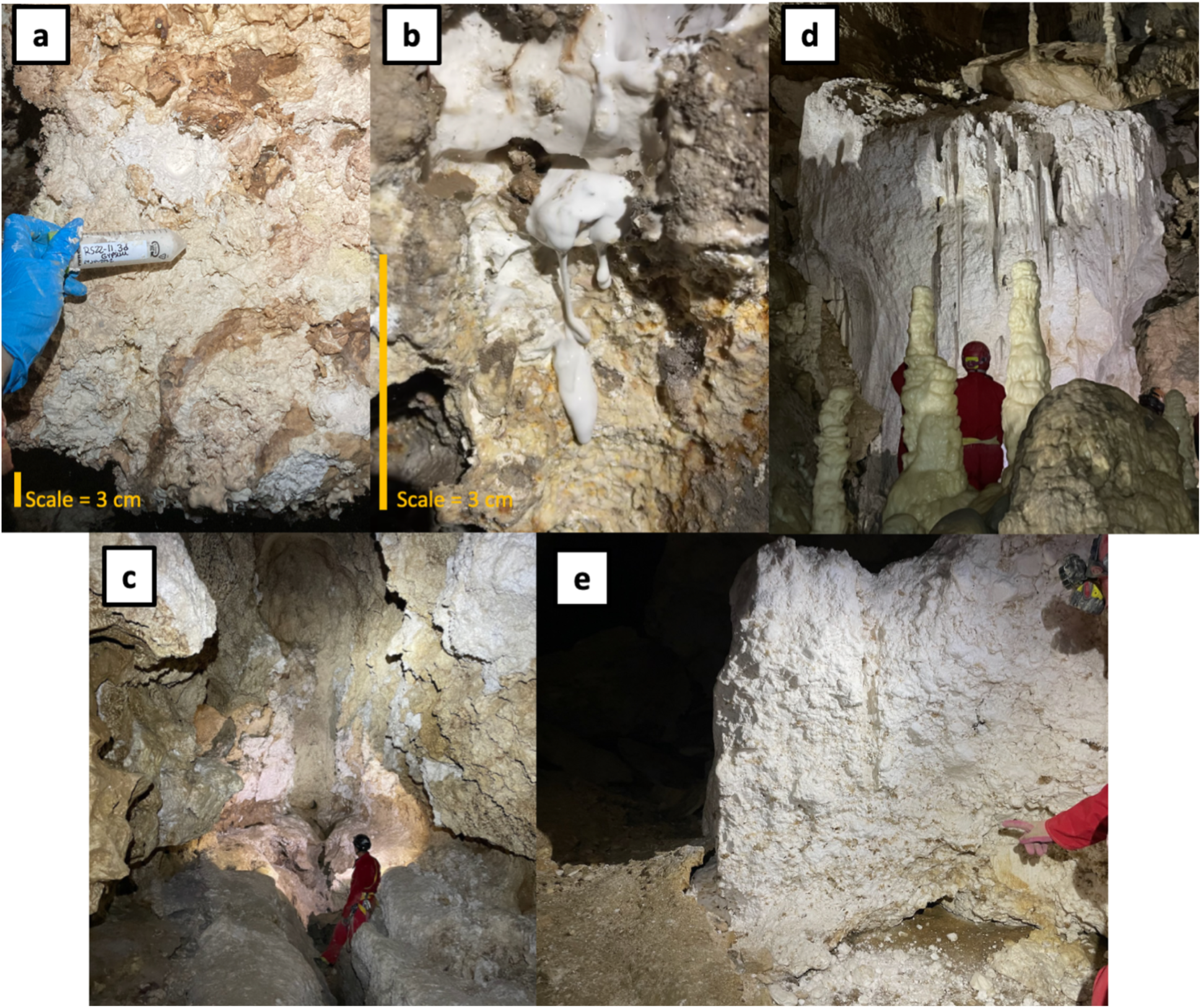
Representative photos of actively forming (photos a-c) and relict gypsum deposits (photos d and e). Panel (a) shows colored crusts that form over the surface of fresh gypsum; (b) shows white freshly-formed gypsum dripping down cave wall; and (c) shows white clumps of microcrystalline gypsum crusts on ceilings 10 meters above a sulfidic stream. Photo (d) is a massive gypsum “glacier” in a relict passage (SDS) from the older levels. Vertical drill hole structures in deposit show where gypsum was dissolved by infiltrating drip waters. Photo (e) is another relict deposit at site TP that has a crumbly and heterogeneous texture. Photos by D. Jones, Z. Havlena, and M. Best.

**Figure 2.**
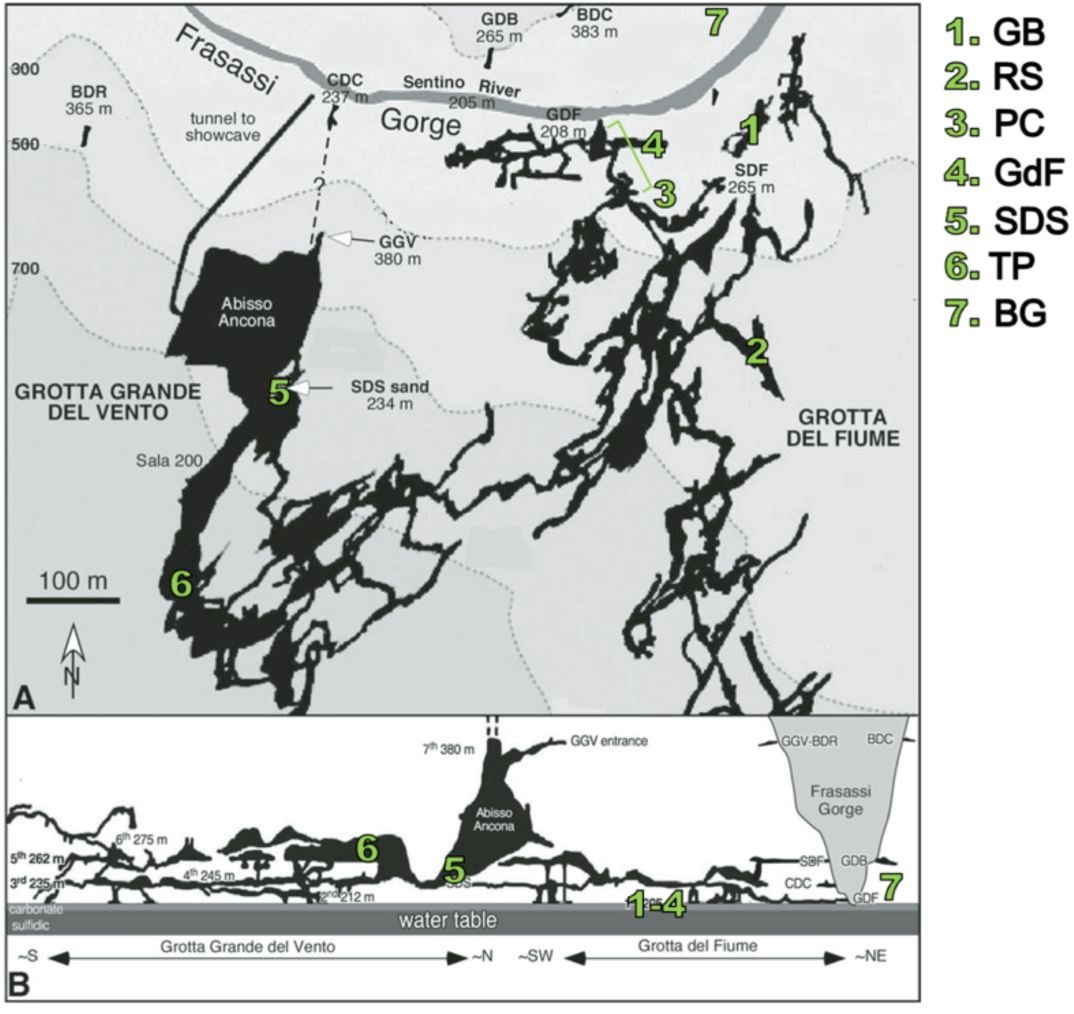
Map of Frasassi cave system, showing sampling locations (Table 1) in map view (A) and in profile (B). Adapted from Montanari et al. (35).

**Table 1.**
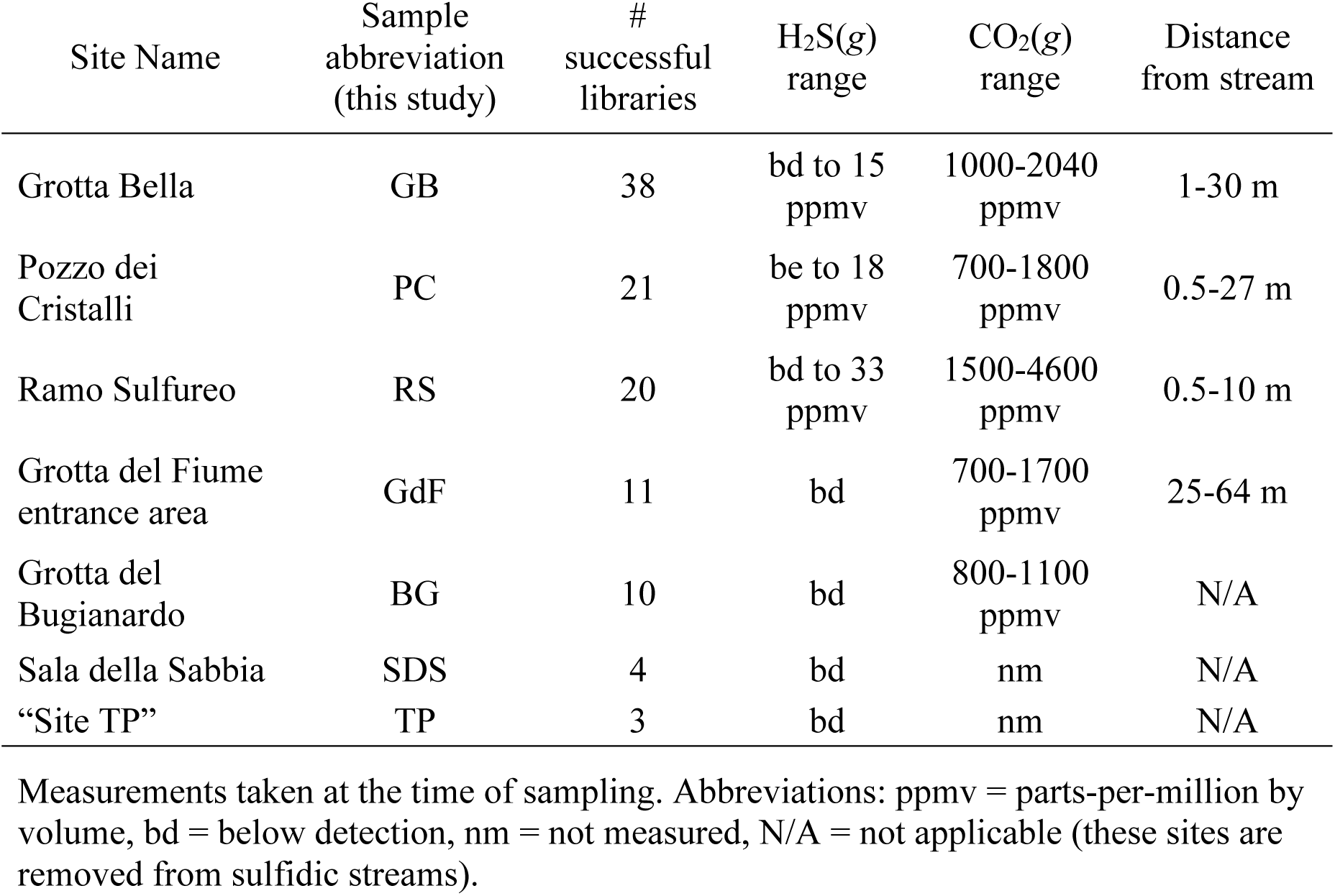
Summary of samples and associated geochemical data.

| Site Name | Sample abbreviation (this study) | # successful libraries | H<sub>2</sub>S(g) range | CO<sub>2</sub>(g) range | Distance from stream |
| --- | --- | --- | --- | --- | --- |
| Grotta Bella | GB | 38 | bd to 15 ppmv | 1000-2040 ppmv | 1-30 m |
| Pozzo dei Cristalli | PC | 21 | be to 18 ppmv | 700-1800 ppmv | 0.5-27 m |
| Ramo Sulfureo | RS | 20 | bd to 33 ppmv | 1500-4600 ppmv | 0.5-10 m |
| Grotta del Fiume entrance area | GdF | 11 | bd | 700-1700 ppmv | 25-64 m |
| Grotta del Bugianardo | BG | 10 | bd | 800-1100 ppmv | N/A |
| Sala della Sabbia | SDS | 4 | bd | nm | N/A |
| “Site TP” | TP | 3 | bd | nm | N/A |

Measurements taken at the time of sampling. Abbreviations: ppmv = parts-per-million by volume, bd = below detection, nm = not measured, N/A = not applicable (these sites are removed from sulfidic streams).

In the older, upper levels, we sampled gypsum from massive breakdown deposits at Sala della Sabbia (SDS) (Fig. 1d) and “Site TP” (TP) on the 5^th^ karst level (Fig. 2) (35), which are accessible by technical caving routes in the show cave portion of Grotta Grande del Vento. These passages are more than 30 m above the current water table and formed more than 100 Ka before present (35, 36). H_2_S was not detectable by smell in these areas. These gypsum deposits were dry and friable. The SDS deposit consists of a lower gray layer with clastic sediment mixed with gypsum, and a massive upper white layer (35), while the deposits at TP were white but more heterogeneous than the white layer at SDS, with larger pink and beige selenite crystals intermixed with powdery white microcrystalline gypsum (Fig. 1e). We also collected samples from Grotta del Bugianardo (site BG), which is a formerly sulfidic cave of unknown age on the opposite side of the Sentino River (42, 43). Gypsum deposits at BG occurred in wall pockets and as breakdown, and were dry, similar to the gypsum from the GdF passage. H_2_S was not detectable in these areas, and CO_2_ concentrations were 800-1100 ppmv.

### Small subunit rRNA gene amplicon libraries

We generated 107 successful 16S rRNA gene libraries, the majority (n= 85) of which were from lower-level passages GB, PC and RS. In hierarchical agglomerative cluster analysis, most samples from these passages separate into three major clusters (Clusters II, III, and IV in Suppl. Fig. S2), whereas most samples from GdF, BG, and the upper deposits from SDS and TP (n=28) cluster together (Cluster I in Suppl. Fig. S2). Libraries in Cluster I therefore appear to represent communities associated with relict gypsum deposits, while libraries in Clusters II-IV represent microbial communities on freshly formed or at least still actively forming gypsum.

Six OTUs were consistently present in Clusters II-IV: *Acidithiobacillus* (OTU 1), *Metallibacterium* (OTU 3), *Mycobacterium* (OTU 6), and three archaea in the *Thermoplasmatales* (OTU 2, *Ferroplasma*; OTU 4, *Cuniculiplasma*; and OTU 5, an uncultured *Thermoplasmatales*). *Acidithiobacillus* is most abundant in Clusters III and IV, while *Metallibacterium* is most abundant in Cluster II and *Mycobacterium* is most abundant in Cluster IV. Among archaea, *Ferroplasma* is most abundant in Cluster IV, while the other two *Thermoplasmatales* OTUs are most abundant in some Cluster II and IV libraries. The representative sequences for OTU 5, the unnamed member of the *Thermoplasmatales*, shares 100% identity with 16S rRNA gene clone RS05_24c_A14 (HM754549) from a Frasassi snottite sample (30), which was classified as a member of the “D-plasma” (44). This OTU is most abundant in Cluster II.

Cluster I was more diverse overall (Suppl. Fig. S3), and is dominated by OTUs from the *Proteobacteria*, with some OTUs from the *Actinomycetota*, *Acidobacteriota*, and *Cyanobacteria*, and includes some unnamed taxa like “Gammaproteobacteria PLTA13” and phylum “WPS-2”. Cluster I also includes multiple process blanks that group together in a smaller cluster within Cluster I, and samples from this subcluster (indicated by a * in Figure S2) were therefore removed from subsequent analyses.

### Relationships between microbial community composition, geochemical variables, and cave location

Libraries from most lower level samples group together in non-metric multidimensional scaling (NMDS) ordinations (Fig. 3a), and were statistically significantly different from clusters of upper level libraries (ANOSIM, R = 0.83, p < 0.001; PERMANOVA, F = 12.9, p < 0.001). When libraries from only Clusters II-IV were ordinated, an environmental overlay shows that distance from the sulfidic stream, as well as concentrations of H_2_S(*g*) and CO_2_(*g*) in the cave atmosphere, were statistically significantly correlated with the ordination (distance: R^2^=0.23, p<0.001; H_2_S: R^2^ = 0.09, p = 0.029; CO_2_: R^2^ = 0.17, p < 0.001). Distance is aligned with the first ordination axes, in the opposite direction from H_2_S(*g*) and CO_2_(*g*) concentrations, consistent with the sulfidic streams as a source of these gases. Species scores for the most abundant taxa show that *Acidithiobacillus* (OTU 1)*, Ferroplasma* (OTU 2), and *Mycobacterium* (OTU 6) are abundant in Clusters III and IV and associated with samples with high H_2_S(*g*) and CO_2_(*g*) concentrations near the sulfidic stream. In contrast, *Metallibacterium* (OTU 3) and the unclassified *Thermoplasmatales* (OTU 5, probably D-plasma) are associated with Cluster III and increased distance from sulfidic streams, while *Cuniculiplasma* (OTU 4) occurs in both Clusters II and IV.

**Figure 3.**
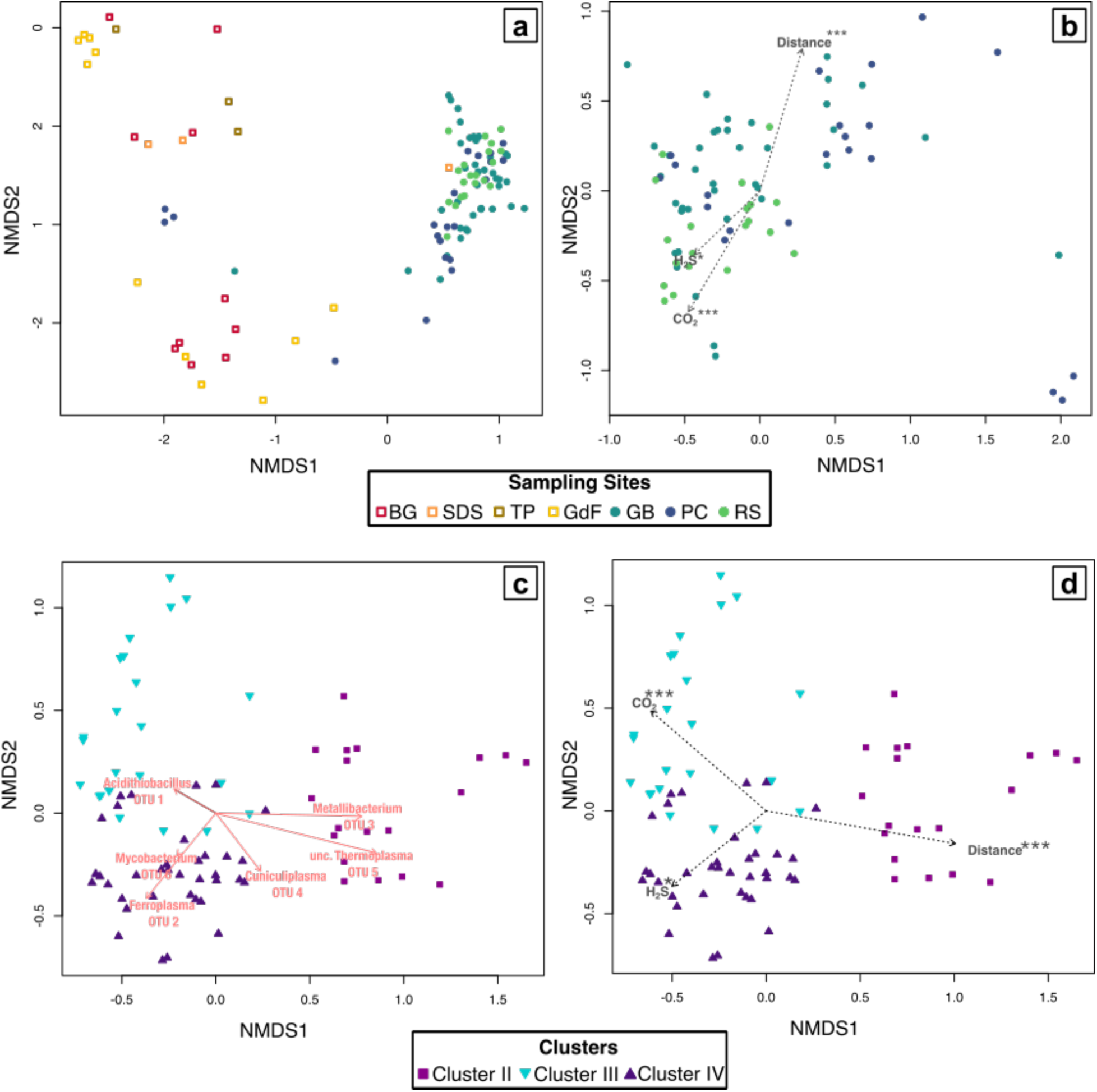
Non-metric multidimensional scaling (NMDS) ordination of rRNA gene libraries (k=4, stress <0.10), comparing (a) libraries from samples collected in the upper level passages (warm colored boxes) and lower level sampling sites (cool colored circles), and (b) libraries from lower level passages alone. Fitted vectors of environmental variables are statistically significantly correlated with the ordination (* = P < 0.05; *** = P < 0.001). Panels (c) and (d) are ordinations of libraries from clusters II-IV in the cluster analysis (Suppl. Fig. 2), interpreted as representing communities from actively-forming gypsum. Panel (c) shows the species scores for the 6 most abundant OTUs (vectors), and panel (d) shows an overlay of fitted vectors of environmental variables.

We therefore examined the specific relationships among these predominant taxa and corresponding variables across all samples from PC, GB, RS, and GdF, for which we were able to measure distance from sulfidic streams. Relative abundance of all *Acidithiobacillus* OTUs decreases with distance from the sulfide source (Pearson’s R = −0.26, p = 0.003; Spearman’s ρ = −0.33, p = 0.003), although this is variable and ranges from >90% to 0% in samples within 10 meters of streams (Fig. 4a). Likewise, *Thermoplasmatales*-group archaea are most abundant close to the stream, but this trend is not statistically significant (Pearson’s R = 0.069, p = 0.53; Spearman’s ρ = 0.09, p = 0.4). Although both *Ferroplasma* and *Cuniculiplasma* occur in samples near the water table, *Ferroplasma* are more abundant in samples with high H_2_S(*g*) concentrations (Table 2), while *Cuniculiplasma* mostly occur in samples with low-undetectable H_2_S(*g*) (Fig. 5).

**Figure 4.**
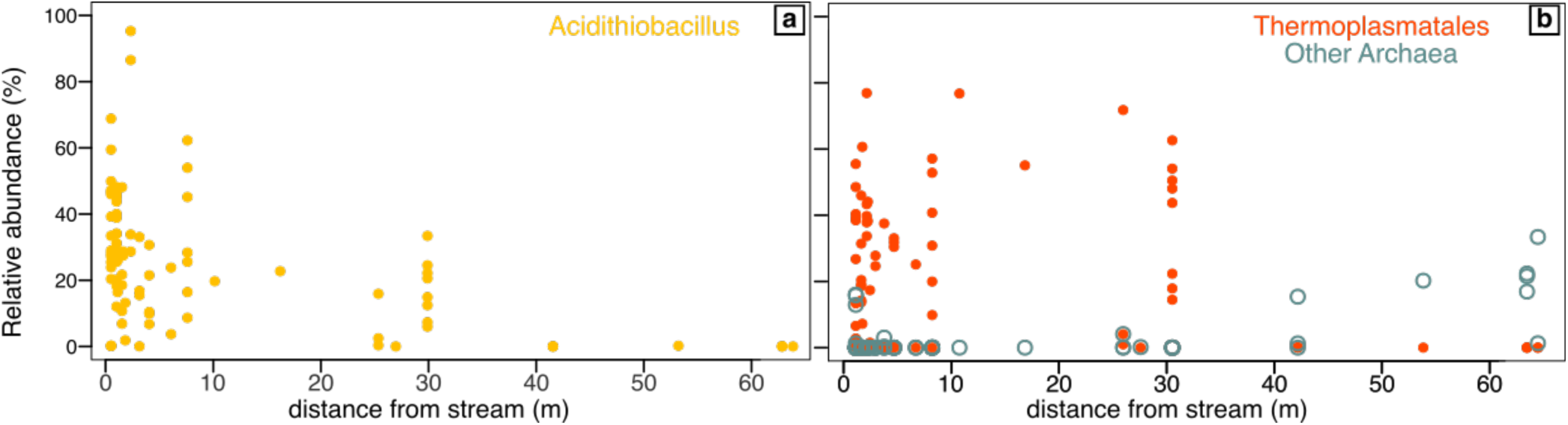
Relative abundance of (a) *Acidithiobacillus* spp. and (b) *Thermoplasmatales* and other archaea in gypsum deposits from samples from the lower levels versus distance from the nearest sulfidic stream.

**Figure 5.**
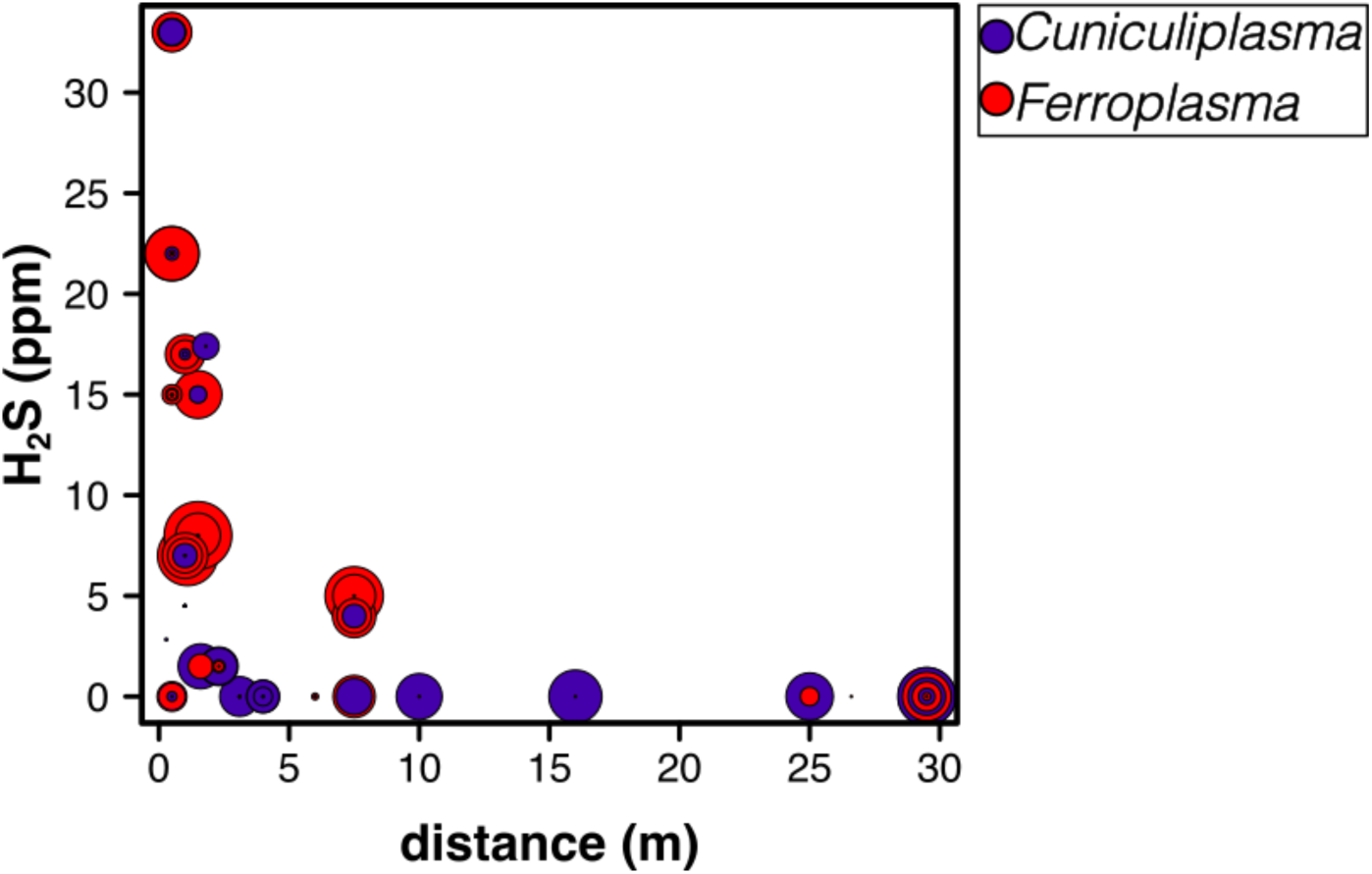
Relative abundance of two most abundant members in *Thermoplasmatales* (*Cuniculiplasma* [OTU 4] and *Ferroplasma* [OTU 2]) compared with H_2_S(*g*) concentration and distance from the nearest sulfidic stream. Points are scaled by relative abundance of each OTU. (Small black dots are samples that do not have significant abundance of either OTU.)

### Cell counts

Cell counts for gypsum surfaces from within 10 m from the sulfidic stream were between 3.4×10^7^-7.2×10^8^ cells g^-1^ (Suppl. Fig. S4). However, further from the stream, cell counts were more variable, as low as 6.3×10^6^ cells g^-1^ to as high as 1.4×10^10^ cells g^-1^. Overall, gypsum surfaces from the older levels had higher cell abundance, between 3.2×10^8^ to 3.3×10^9^ cells g^-1^ (Suppl. Fig. S4).

## Discussion

### Microbial communities from actively-forming gypsum deposits

The first goal of this study was to characterize microbial communities in actively forming cave gypsum, and evaluate how communities change further from the sulfidic water table. While many studies have focused on snottite biofilms (25, 31, 45, 46), to our knowledge, the only study that has described microbial communities on freshly-formed SAS gypsum was from a seawater-influenced sulfidic cave by D’Angeli et al. (47). D’Angeli et al. (47) found abundant *Thermoplasma* and other archaea in the *Thermoplasmatales* group, with lower amounts of bacterial taxa such as *Acidithiobacillus*, *Metallibacterium*, and *Sulfobacillus*, some of which are also known from sulfidic cave snottites (30, 31). Accordingly, we expected that microbial communities from actively-forming gypsum in Frasassi would contain many of the same organisms that are known from Frasassi snottites, and that these communities would change with distance from degassing sulfidic streams where H_2_S(*g*) flux is lower and, therefore, less chemical energy is available.

The taxa that were consistently abundant in fresh gypsum at all lower-level passages (RS, PC, and GB) were *Acidithiobacillus*, *Metallibacterium*, *Mycobacterium,* and Thermoplasmatales-group archaea. We interpret this microbial assemblage as indicative of the community associated with actively forming SAS gypsum. Based on hierarchical agglomerative cluster analysis and NMDS ordinations (Fig. 3 and Suppl. Fig. 2), the gypsum community closest to the water table where H_2_S(*g*) flux is highest was dominated by *Acidithiobacillus*, *Ferroplasma*, and *Mycobacterium*. Gypsum that was further away (but still actively forming in the lower levels) had more *Metallibacterium* and other *Thermoplasmatales*. There seems to be niche separation among the extremely acidophilic archaea, with *Ferroplasma* consistently more abundant in samples exposed to high H_2_S(*g*) concentrations, while *Cuniculiplasma* occurs in samples with low H_2_S(*g*) and that are further from the stream (Fig. 5). Based on metagenomic analysis (46), *Cuniculiplasma* from Frasassi have a sulfide-quinone reductase but do not appear to encode other enzymes for the oxidation of reduced inorganic sulfur compounds, while *Ferroplasma* from sulfidic caves have a more complete sulfide oxidation pathway including SQR and sulfur-oxygenase reductase (SOR). Likewise, *Acidithiobacillus thiooxidans* are well-known sulfide oxidizers (48), and *Metallibacterium* are also capable of oxidizing some reduced inorganic sulfur compounds (49, 50). Therefore, four of the most abundant members of the gypsum-associated communities may be capable of sulfide oxidation (*Acidithiobacillus*, *Metallibacterium*, *Ferroplasma*, and *Cuniculiplasma*) and may therefore be directly contributing to sulfuric acid production and gypsum formation in the cave. As far as the other two most abundant members, no genomic data is available for D-plasma (to our knowledge), and *Mycobacterium* spp. have also been found on subaerial surfaces in other sulfur-rich cave surfaces, where they appear to be methanotrophs or organotrophs (51, 52).

Many of the organisms on gypsum surfaces, particularly *Acidithiobacillus*, *Ferroplasma*, and *Cuniculiplasma* (formerly G-plasma), are also abundant in snottite biofilms from the same cave (30, 31, 33). However, *Acidithiobacillus* is much more abundant in snottites than in gypsum. Based on fluorescence *in situ* hybridization of 16 snottite samples from Frasassi, *Acidithiobacillus* averaged 68.6% of total cells, and were sometimes the only microorganisms detected, while archaea averaged 18.2% of total cells and were at most 38.6% (30). In contrast, here we found that *Acidithiobacillus* averaged 29.6% of 16S rRNA genes in actively-forming gypsum, and Thermoplasmatales group archaea were 19.8%. This higher proportion of *Acidithiobacillus* spp. in snottites is consistent with indications that *Acidithiobacillus* may be the primary snottite-forming organism (45), and is therefore more abundant in biofilms than gypsum surfaces.

DNA extraction was challenging from the cave gypsum. In previous experiments with Frasassi gypsum, Havlena et al. (53) showed that dissolving gypsum prior to extraction did not improve DNA recovery. Overall, gypsum has far fewer cells than other cave deposits such as biovermiculations, which have up to 10^11^ cells g^-1^ (54), so low DNA recovery from gypsum may have been a result of low microbial biomass. In addition, following sample collection, these molecules could be rapidly degrading in the oxidizing, acidic conditions of the fresh gypsum (55).

### Relict gypsum surfaces

The second goal of this study was to evaluate if and how microbes continue to use gypsum as a habitat in older, relict gypsum deposits. Microbial communities from massive gypsum breakdown deposits in the upper levels were distinct from those on most deposits in the lower, active levels (Fig. 3). All successful libraries from GdF, BG, and the upper older levels (SDS and TP) clustered together with some libraries from deposits that were far from the water table in PC and GB (Cluster I in Suppl. Fig. 2). These libraries do not contain more than 1% of the chemolithotrophic taxa associated with fresh gypsum, and instead contain more diverse microbial communities with taxa that are likely organotrophs such as *Brevibacillus* and *Lactobacillus*, as well as other populations such as the gammaproteobacterial taxon “wb1-P19” that is a methanotroph (56) and possible inorganic nitrogen oxidizers such as the crenarchaeon “*Candidatus* Nitrosotalea” (57). Surprisingly, however, they have higher cell numbers than freshly forming gypsum (Fig. S4), which perhaps reflects the less acidic mineral matrix of the older deposits. In addition, the relict gypsum deposits are presumably less dynamic than freshly-forming gypsum, so cells may have had more time to accumulate on the surfaces of the older deposits. Future work should address whether cells on these older gypsum deposits are active, dormant, or dead.

These distinct microbial assemblages likely reflect different energy resources for gypsum surface communities. The upper levels of Frasassi are not currently exposed to H_2_S(*g*) from the sulfidic water table, and gypsum only remains in places that are protected from dripping surface waters, so microorganisms on gypsum surfaces must rely on trace gases or particulate material transported in air currents. Frasassi is also one of Italy’s most popular show caves, with more than 250,00 visitors to the upper levels each year (58), so material from tourists likely influences these communities. However, libraries from sites SDS and TP in the show cave portion were similar to those from relict gypsum deposits in sites BG, GdF, PC, and GB, which are not part of the show cave and some of which are only accessible by technical caving routes. These communities may therefore represent an organotroph-dominated community that typifies relict gypsum in ancient sulfuric acid caves.

### Astrobiological implications

Exploring these gypsum-associated communities and their distribution enhances our understanding of how life might develop and be detected on other planetary bodies with similar salty and acidic conditions. The Frasassi Caves are an exceptional setting in which to explore subsurface microbial life adapted to acidic, sulfate-precipitating environments, both while the gypsum is freshly forming but also in the relict deposits in the upper cave levels that have dried out over time. Furthermore, because of its subterranean location, these deposits do not contain photosynthetic life and are isolated from organic material from the surface (59), so the extremophilic communities in Frasassi offer a potential window into worlds in which photosynthesis never evolved (2). We showed here that freshly-forming gypsum in Frasassi is colonized by low biomass communities of chemolithotrophic bacteria and archaea that take advantage of inorganic chemical energy available in the cave atmosphere, and seem to be organized in niches based on energy availability that reflects the spatial variability of energy resources within the cave ecosystem (Figs. 4 and 5). In contract, older deposits have a different microbial community that is adapted to lower energy conditions, and may have overprinted the original community.

We don’t know if and how faithfully SAS gypsum preserves biosignatures of the chemosynthetic communities that are associated with freshly forming gypsum. Actively forming gypsum in Frasassi contains isotopic signatures of microbial H_2_S oxidation (29), and it and other sulfidic caves contain other potential morphological, mineral, and isotopic biosignatures (60, 61). Studies have also showed that gypsum deposits in other acidic environments contain morphological biosignatures and intact cells (4, 22). Terrestrial analog sites are most robust where multiple lines of biosignatures are present, as is necessary for any extraplanetary setting (2). However, we don’t know if these deposits preserve molecular biosignatures over longer periods of time, and so far, organic biomarkers associated with sulfidic cave gypsum have not been determined. Several of the most abundant taxa that we identified on actively-forming gypsum are capable of producing lipid biomarkers that are valuable for astrobiology studies (62). *Acidithiobacillus* spp. and *Thermoplasmatales* from cave snottites produce bacteriohopanepolyols (BHPs) and glycerol dialkyl glycerol tetraethers (GDGTs), respectively (45, 63), and we assume that the *Acidithiobacillus* and archaea identified here do as well. Based on the abundance of *Acidithiobacillus* and *Thermoplasmatales*, either of these molecule classes would be valuable for evaluating organic biomarker preservation in acidic gypsum. However, the presence of distinct communities dominated by organotrophic communities on Frasassi’s relict gypsum also suggests that the initial gypsum-precipitating consortia might be overprinted if the biomarkers are not trapped within breakdown deposits.

## Methods

### Sample collection

Gypsum was sampled using sterile implements, usually by collecting triplicate samples within 50 cm from the same wall location. Aliquots for DNA extraction were preserved in RNAlater (ThermoFisher Scientific, Waltham, MA, USA), and aliquots for microscopy were fixed within 12-16 hours of collection in 4% PFA following the procedure of Jones et al. (2023). The pH of damp gypsum in low level sites was recorded using test strips.

At the time of sampling, concentrations of H_2_S, O_2_, SO_2_, NH_3_, CO, LEL, and CO_2_ were recorded with handheld meters (RECON/4a and Sapphire, ENMET, Ann Arbor, MI, USA), and sometimes for H_2_S, CO_2_, and SO_2_, Dräger tubes (Draegerwerk AG & Co., Lübeck Germany). H_2_S(*g*) interferes with the NH_3_ detector (Sensoric NH3 3E 100 SE, Honeywell, Charlotte, NC, USA) on the ENMET Sapphire over 20 ppmv, so we only report data from areas where H_2_S is less than 10 ppmv. We also measured distance from the nearest cave stream as a proxy for influence of gases from the sulfidic water table. Humidity and temperature at these locations have been measured previously and are stable year-round at nearly 100% and ∼13°C (29).

### Microscopy and cell counts

Direct DAPI imaging was performed as described in (53). Cell counting was performed as in (64), in which PFA-fixed aliquots were sonicated for 45 seconds using a Fisherbrand model 120 dismembrator (Thermofisher). Water content of the gypsum was determined by drying small aliquots at 60°C and recording the weight before and after. Cells abundance was reported as cells per gram of dry sediment.

### DNA extraction and rRNA gene library preparation

DNA extraction and rRNA gene libraries were prepared using the procedure described in detail in Havlena et al. (53). Briefly, DNA was extracted with a DNeasy Powersoil Pro DNA extraction kit (Qiagen, Germantown, MD, USA), using a bead-beating protocol selected for the best yield (53). RNAlater was removed prior to extraction by diluting 1:1 with PCR water or molecular grade PBS, centrifuging, and removing the supernatant. Libraries were prepared using Nextera amended primers 515f and 806r (f: TCG TCG GCA GCG TCA GAT GTG TAT AAG AGA CAG; r: GTC TCG TGG GCT CGG AGA TGT GTA TAA GAG ACA G), amplified with HotStarTaq Plus (Qiagen) polymerase with 5 min for initial denaturation (94°C), 35 cycles with 45 s denaturation (94°C), 60 s annealing (50°C), 90 s elongation (72°C), and 10 min final elongation (72°C). Process blanks (negative controls) were included during extraction and PCR steps, and submitted along with sample extracts to the University of Minnesota Genomics Center (UMGC) for barcoding and sequencing using on an Illumina MiSeq (300 bp paired-end cycles).

## Data availability

rRNA gene libraries from this study were deposited under BioProject accession number (PRJNA1220587) in the Sequence Read Archive (SRA; http://www.ncbi.nlm.nih.gov/sra).

## Bioinformatics and statistical analysis

Processing of high-throughput 16S rRNA gene libraries followed the same approach outlined in Havlena et al. (53). Raw fastq-formatted libraries were processed by trimming and removing adapters with Sickle v1.33 (https://github.com/najoshi/sickle) and cutadapt (65), respectively. Read merging, dereplication, and clustering into operational taxonomic units (OTUs, 97% similarity) were performed with PEAR (66), VSEARCH v2.21 (67), and the UPARSE pipeline (68) as in (53). Taxonomic classifications (confidence score ≥50) were assigned using the Silva v138 database (69) in mothur v 1.36.1 (70).

Statistical analyses were performed in R v. 4.2.3 (71) using the vegan (v. 2.6-2, (72)) and cluster (v. 2.1.6, (73)) packages. OTUs were analyzed after libraries with <10,000 total sequence reads were removed from the dataset, and then raw OTU counts were transformed to proportion of total for remaining libraries. For cluster and ordination analyses, the proportion matrix was additionally transformed using Hellinger transformation 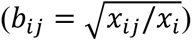 where *x_i,j_* is an element in the matrix of proportional values and *b_i,j_* is an element in the square-root transformed matrix. Non-metric multidimensional scaling (NMDS) ordinations were preformed using the *metaMDS* function, with Bray-Curtis distance and 4 dimensions (k=4). Hierarchical agglomerative clustering used Euclidian distance and Ward’s linkage method. Clustering was performed for Q-mode (samples, all OTUs >0.01% of total) and R-mode (top 50 most abundant OTUs).

Calculations of difference among samples at variable levels used ANOSIM and PERMANOVA functions in vegan (default parameters), and pairwise post-hoc analysis with *pairwiseadonis* function (74).

## Acknowledgements

We thank A. Montanari and P. Metallo for logistical support and the use of facilities and laboratory space at the Osservatorio Geologico di Coldigioco. Special thanks to M. Best, S. Recanatini, S. Mariani, S. Cerioni, I. D’Angeli, I. Vaccarelli, H. Aronson, S. Morin, and J. Macalady for technical support and assistance with fieldwork and sample collection. The staff at the University of Minnesota Genomics Center provided insightful discussion and assistance with amplicon sequencing of challenging samples. This research was primarily supported by the NASA Exobiology program (80NSSC20K0619), the NASA Future Investigators in NASA Earth and Space Science and Technology (FINESST) program (80NSSC21K1547) to DSJ and ZEH, with additional support from New Mexico Institute of Mining and Technology and the National Cave and Karst Research Institute.

