## Supplementary Figures for "Microbial ecology of acidic, biogenic gypsum: Community structure and distribution of extremophiles on freshly formed and relict sulfate deposits in a hydrogen sulfide-rich cave"

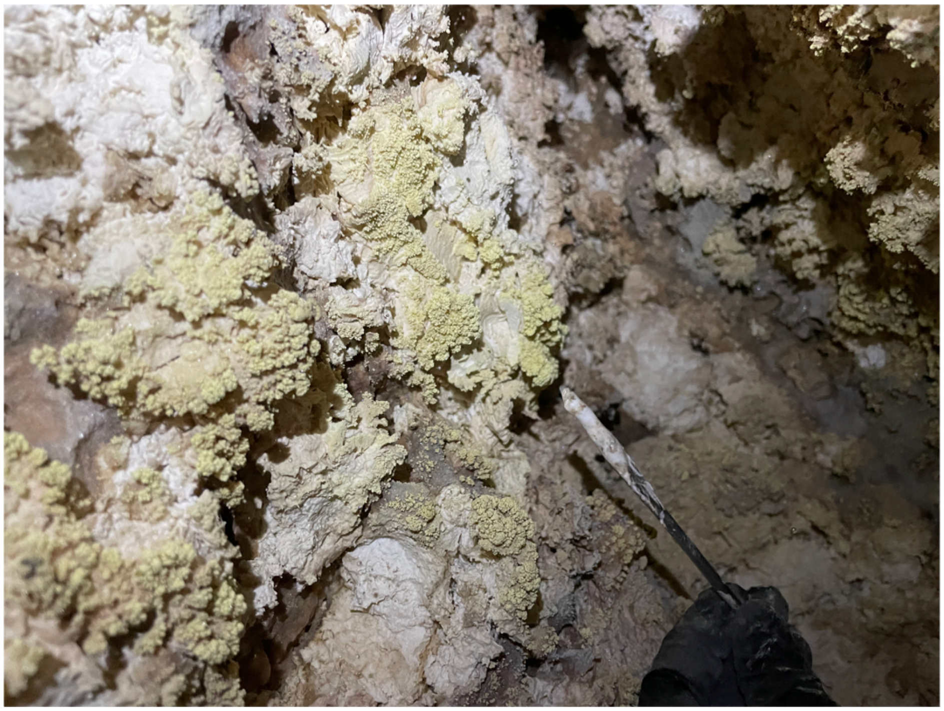

**Supplemental Figure S1.** Photo from gypsum at site RS, where elemental sulfur (S°) is present as yellow coatings on or near white microcrystalline gypsum.

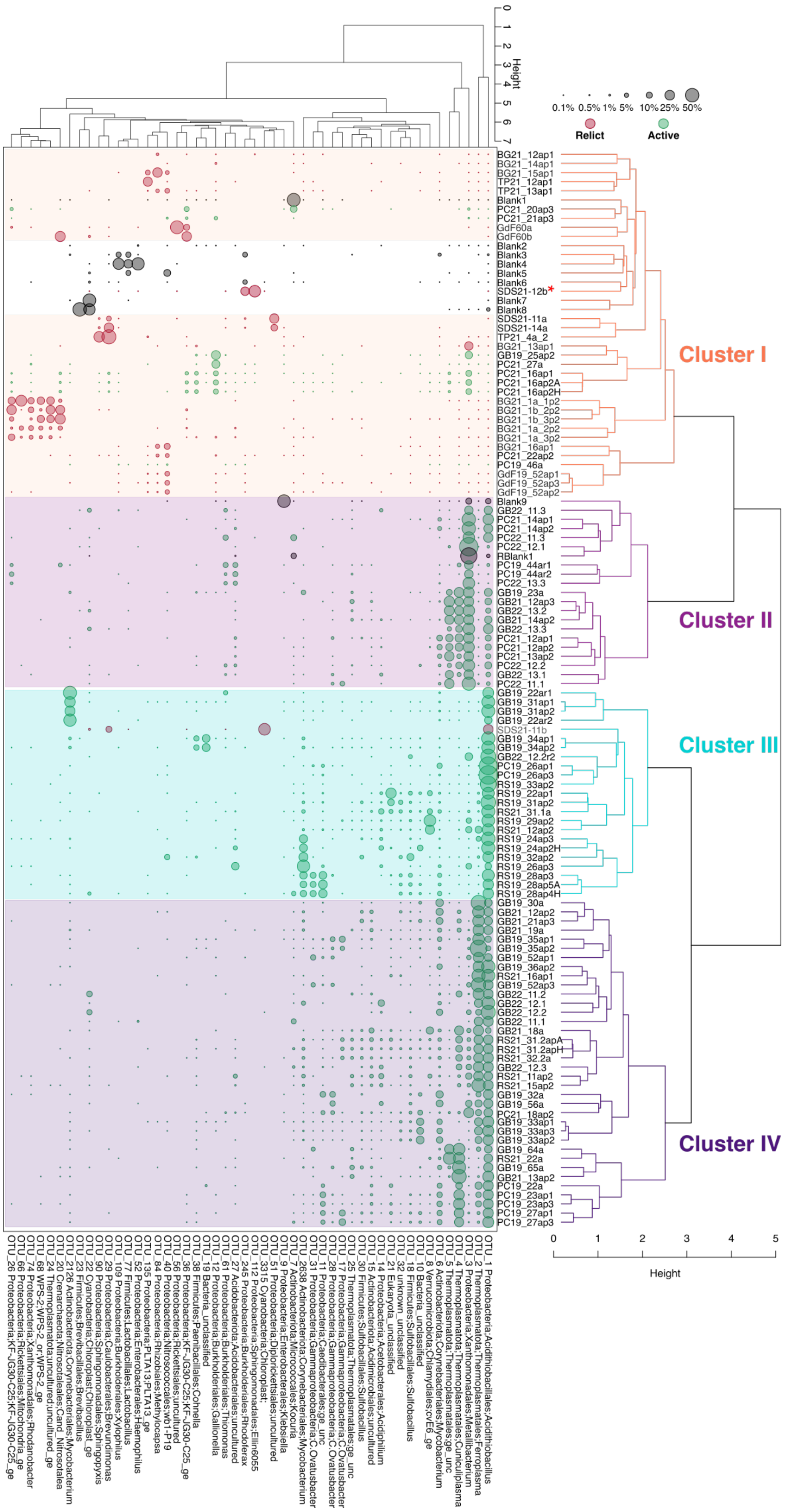

Height
0
1
2
3
4
5
6
7
0.1% 0.5% 1% 5% 10% 25% 50%
Relict
Active
BG21_12ap1
BG21_14ap1
BG21_15ap1
TP21_12ap1
TP21_13ap1
Blank1
PC21_20ap3
PC21_21ap3
GdF60a
GdF60b
Blank2
Blank3
Blank4
Blank5
Blank6
SDS21-12b*
Blank7
Blank8
SDS21-11a
SDS21-14a
TP21_4a_2
BG21_13ap1
GB19_25ap2
PC21_27a
PC21_16ap1
PC21_16ap2A
PC21_16ap2H
BG21_1a_1p2
BG21_1b_2p2
BG21_1b_3p2
BG21_1a_2p2
BG21_1a_3p2
BG21_16ap1
PC21_22ap2
PC19_46a
GdF19_52ap1
GdF19_52ap3
GdF19_52ap2
Blank9
GB22_11.3
PC21_14ap1
PC21_14ap2
PC22_11.3
PC22_12.1
RBlank1
PC19_44ar1
PC19_44ar2
PC22_13.3
GB19_23a
GB21_12ap3
GB22_13.2
GB21_14ap2
GB22_13.3
PC21_12ap1
PC21_12ap2
PC21_13ap2
PC22_12.2
GB22_13.1
PC22_11.1
GB19_22ar1
GB19_31ap1
GB19_31ap2
GB19_22ar2
SDS21-11b
GB19_34ap1
GB19_34ap2
GB22_12.2r2
PC19_26ap1
PC19_26ap3
RS19_33ap2
RS19_22ap1
RS19_31ap2
RS21_31.1a
RS19_29ap2
RS21_12ap2
RS19_24ap3
RS19_24ap2H
RS19_32ap2
RS19_26ap3
RS19_28ap3
RS19_28ap5A
RS19_28ap4H
GB19_30a
GB21_12ap2
GB21_21ap3
GB21_19a
GB19_35ap1
GB19_35ap2
GB19_52ap1
GB19_36ap2
RS21_16ap1
GB19_52ap3
GB22_11.2
GB22_12.1
GB22_12.2
GB22_11.1
GB21_18a
RS21_31.2apA
RS21_31.2apH
RS21_32.2a
GB22_12.3
RS21_11ap2
RS21_15ap2
GB19_32a
GB19_56a
PC21_18ap2
GB19_33ap1
GB19_33ap3
GB19_33ap2
GB19_64a
RS21_22a
GB19_65a
GB21_13ap2
PC19_22a
PC19_23ap1
PC19_23ap3
PC19_27ap1
PC19_27ap3
Cluster I
Cluster II
Cluster III
Cluster IV
0
1
2
3
4
5
Height
OTU_1 Proteobacteria;Acidithiobacillales;Acidithiobacillus
OTU_2 Thermoplasmatota;Thermoplasmatales;Ferroplasma
OTU_3 Proteobacteria;Xanthomonadales;Metallibacterium
OTU_4 Thermoplasmatota;Thermoplasmatales;Cuniculiplasma
OTU_5 Thermoplasmatota;Thermoplasmatales;ge_unc
OTU_6 Actinobacteriota;Corynebacteriales;Mycobacterium
OTU_8 Verrucomicrobiota;Chlamydiales;cvE6_ge
OTU_10 Bacteria_unclassified
OTU_18 Firmicutes;Sulfobacillales;Sulfobacillus
OTU_32 unknown_unclassified
OTU_21 Eukaryota_unclassified
OTU_14 Proteobacteria;Acetobacterales;Acidiphilium
OTU_15 Actinobacteriota;Acidimicrobiales;uncultured
OTU_30 Firmicutes;Sulfobacillales;Sulfobacillus
OTU_25 Thermoplasmatota;Thermoplasmatales;ge_unc
OTU_17 Proteobacteria;Gammaproteobacteria;C.Ovatusbacter
OTU_28 Proteobacteria;Gammaproteobacteria;C.Ovatusbacter
OTU_11 Proteobacteria;Caedibacterales;ge_unc
OTU_31 Proteobacteria;Gammaproteobacteria;C.Ovatusbacter
OTU_2638 Actinobacteriota;Corynebacteriales;Mycobacterium
OTU_7 Actinobacteriota;Micrococcales;Kocuria
OTU_9 Proteobacteria;Enterobacterales;Klebsiella
OTU_51 Proteobacteria;Diplorickettsiales;uncultured
OTU_3315 Cyanobacteria;Chloroplast;
OTU_112 Proteobacteria;Sphingomonadales;Ellin6055
OTU_245 Proteobacteria;Burkholderiales;Rhodoferax
OTU_27 Acidobacteriota;Acidobacteriales;uncultured
OTU_61 Proteobacteria;Burkholderiales;Thiomonas
OTU_12 Proteobacteria;Burkholderiales;Gallionella
OTU_19 Bacteria_unclassified
OTU_38 Firmicutes;Paenibacillales;Cohnella
OTU_36 Proteobacteria;KF-JG30-C25;KF-JG30-C25_ge
OTU_56 Proteobacteria;Rickettsiales;uncultured
OTU_40 Proteobacteria;Nitrosococcales;wb1-P19
OTU_84 Proteobacteria;Rhizobiales;Methylocapsa
OTU_135 Proteobacteria;PLTA13;PLTA13_ge
OTU_52 Proteobacteria;Enterobacterales;Haemophilus
OTU_77 Firmicutes;Lactobacillales;Lactobacillus
OTU_109 Proteobacteria;Burkholderiales;Xylophilus
OTU_29 Proteobacteria;Caulobacterales;Brevundimonas
OTU_90 Proteobacteria;Sphingomonadales;Sphingopyxis
OTU_22 Cyanobacteria;Chloroplast;Chloroplast_ge
OTU_23 Firmicutes;Brevibacillales;Brevibacillus
OTU_2126 Actinobacteriota;Corynebacteriales;Mycobacterium
OTU_20 Crenarchaeota;Nitrosotaleales;Cand_Nitrosotalea
OTU_24 Thermoplasmatota;uncultured;uncultured_ge
OTU_68 WPS-2;WPS-2_or;WPS-2_ge
OTU_74 Proteobacteria;Xanthomonadales;Rhodanobacter
OTU_66 Proteobacteria;Rickettsiales;Mitochondria_ge
OTU_26 Proteobacteria;KF-JG30-C25;KF-JG30-C25_ge

**Supplemental Figure S2 (above).** Two-way cluster analysis of 16S rRNA gene libraries. Samples were clustered (Q-mode cluster analysis) using all OTUs, while OTUs were clustered (R-mode cluster analysis) using the 48 most abundant OTUs. Relative abundance of OTUs is shown with dots that are scaled to the proportion of OTUs in each library. The unshaded portion of Cluster I indicates the subcluster associated with process blanks, where the asterisk indicates gypsum sample excluded from downstream analyses.

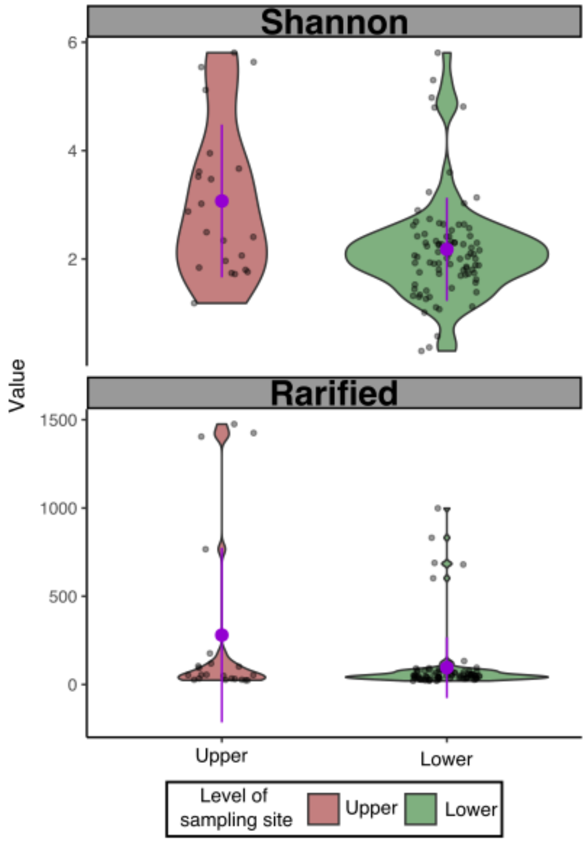


**Supplemental Figure S3.** Alpha diversity of samples from upper versus lower levels. The top panel shows Shannon diversity, and the bottom panels shows richness, both after rarefying libraries to 10,000 sequences. The purple dot and error bars indicate mean and standard deviation.

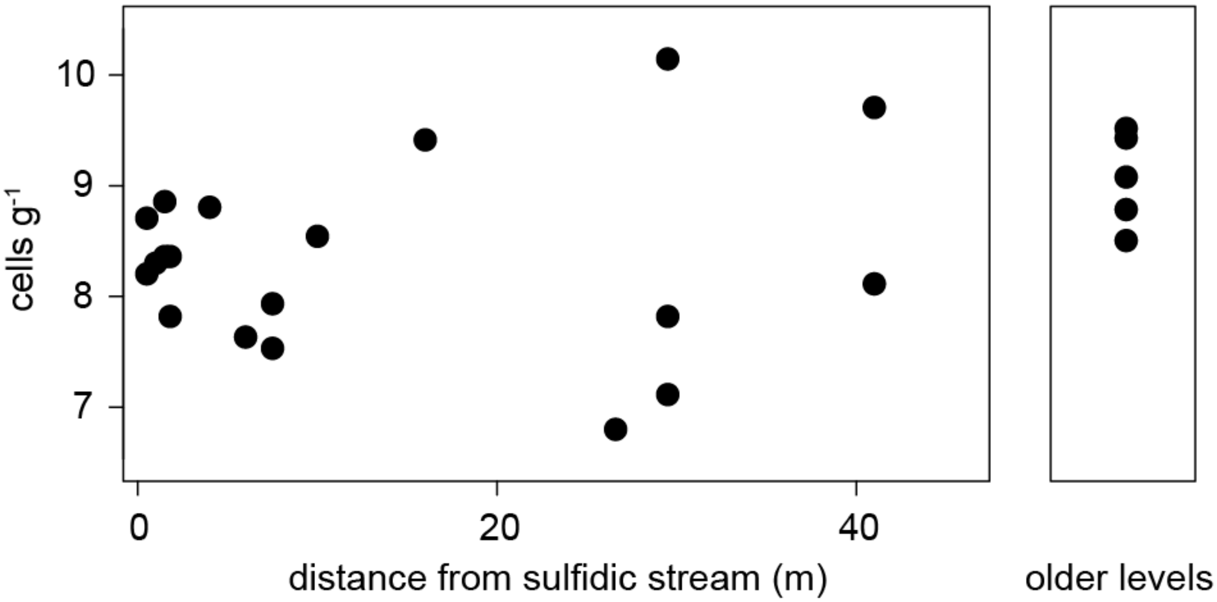


**Supplemental Figure S4.** Cell counts from gypsum surfaces.
